# Effects of sexual dimorphism and estrous cycle on *C. difficile* infections prophylaxis in two rodent models

**DOI:** 10.1101/2025.05.21.655361

**Authors:** Jacqueline R. Phan, McKenzie Washington, Dung M. Do, Tiffany V. Mata, Maria Niamba, Efren Heredia, Robert Soriano, Chandler Hassan, Chad L. Cross, Ernesto Abel-Santos

**Affiliations:** Department of Chemistry and Biochemistry, University of Nevada, Las Vegas, Las Vegas, NV, 89154; Department of Physical and Life Sciences, Nevada State College, Henderson, NV 89002; Department of Epidemiology & Biostatistics, School of Public Health, University of Nevada, Las Vegas, Las Vegas, NV, 89154

## Abstract

*Clostridioides difficile* infection (CDI) is responsible for the majority of identifiable hospital-related antibiotic-associated diarrhea. Susceptibility to CDI and severity of disease varies depending on a variety of factors such as aggressive use of broad-spectrum antibiotics, age, and immune status. Epidemiological studies have consistently shown that female patients are more at risk for CDI than their male counterparts. *C. difficile* is spread by spores which can persist in the environment and in the intestines of patients. Spores do not cause disease but germinate in the antibiotic-altered gut of patients to generate toxin producing vegetative cells. The germination of *C. difficile* spores is mediated by the composition of bile salts in the gut with taurocholate facilitating germination and chenodeoxycholate inhibiting it.

---

Since germination is required for CDI, we expect that anti-germination therapy will prevent CDI and its relapse. While the mechanism for germination has not been unambiguously identified, we and others have shown that *C. difficile* spore germination can be inhibited with bile salt analogs. We also found that cholan-24-amides containing *m*-sulfanilic acid (CamSA) or aniline (CaPA) prevented CDI in rodents.

In this study, we show that prophylactic treatment of murine CDI also showed sexual dimorphism with females responding better to treatment than males. Interestingly, infection sexual dimorphism was reversed in hamsters, with male hamsters developing more severe CDI signs than females. Furthermore, female mice responded better to CDI prophylaxis than their male counterparts.

## Background

*Clostridioides difficile*, the etiological agent of *C. difficile* infection (CDI), is an anaerobic, spore-forming bacterial that is responsible for approximately 25% of all antibiotic-associated diarrhea [1]. CDI has been designated as an urgent threat by the CDC [2]. In the U.S., over 223,900 CDI cases occur annually, with a mortality rate of up to 6.5% and costs estimated to be $6.3 billion [3]. The incidence of CDI is complicated by the appearance of highly resistant and hypervirulent strains [4, 5].

CDI treatments are limited. Recent guidelines discourage the use of cheap metronidazole due to the propensity for triggering relapses [6]. Oral vancomycin is now recommended for primary CDI [7]. Fidaxomicin is a newer antibiotic that causes fewer CDI relapses, but it is significantly more costly [8, 9]. Monoclonal antitoxins can reduce CDI morbidity and mortality, but do not reduce *C. difficile* colonization [10, 11]. Fecal transplantation has been used effectively as CDI treatment [12] but can potentially lead to transfer of deadly pathogens [13]. Despite the high initial response rate to therapy, recurrence of symptoms develops in up to 20% of patients. CDI relapse is due to *C. difficile* spores persisting around the environment of the patient with subsequent germination in the intestine. Therapy for multiple CDI relapses of *C. difficile* colitis has not been examined by randomized, prospective, controlled clinical trials and the best therapeutic approach is currently uncertain [7].

Epidemiological studies have shown that antibiotic use is the main risk factor associated with CDI [14, 15]. Immunocompetence is also strongly correlated with CDI symptom onset [16–18]. Furthermore, older populations are more susceptible to CDI than younger cohorts [14].

When adjusted for age, immune status, and antibiotic exposure, sex is an important risk factor in human CDI. Contrary to other GI bacterial infections [19], women are more vulnerable than men to both primary CDI and CDI relapse [20–27]. More recently, CDI has been flagged as an emerging threat to pregnant and peripartum women [28–31]. Similarly, older women might be at higher risk of developing CDI.

An obvious factor underlying human sexual dimorphisms is the fluctuating hormone concentrations during the interdependent ovarian cycle and uterine (menstrual) cycle [32]. Similar to humans, the estrous cycle of female rodents passes through four different stages: proestrus, estrus, metestrus, and diestrus, [33]. Transitions between estrous stages are associated with sex hormones fluctuations and uterine cellular changes [34].

*C. difficile* forms spores that act as infection vehicles for CDI [35]. Spores do not cause disease but recognize bile salts to germinate into toxin-producing cells [36–41]. Since *C. difficile* spore germination is a required first step in establishing infection in both humans and animals, we developed anti-germination approaches to provide a unique prophylactic therapy for human CDI [42]. With this goal, have examined bile salt and steroidal hormones as competitive inhibitors of spore germination [43, 44]. We found that the amide-linked cholates CamSA and CaPA (**Fig. 1**), inhibit *C. difficile* spore germination at micromolar concentrations even when the taurocholate germinant is present at saturating millimolar levels [45]. As our first CDI prophylactics, CamSA and CaPA have been extensively characterized *in vitro* and *in vivo* [46–52].

**Figure 1.**
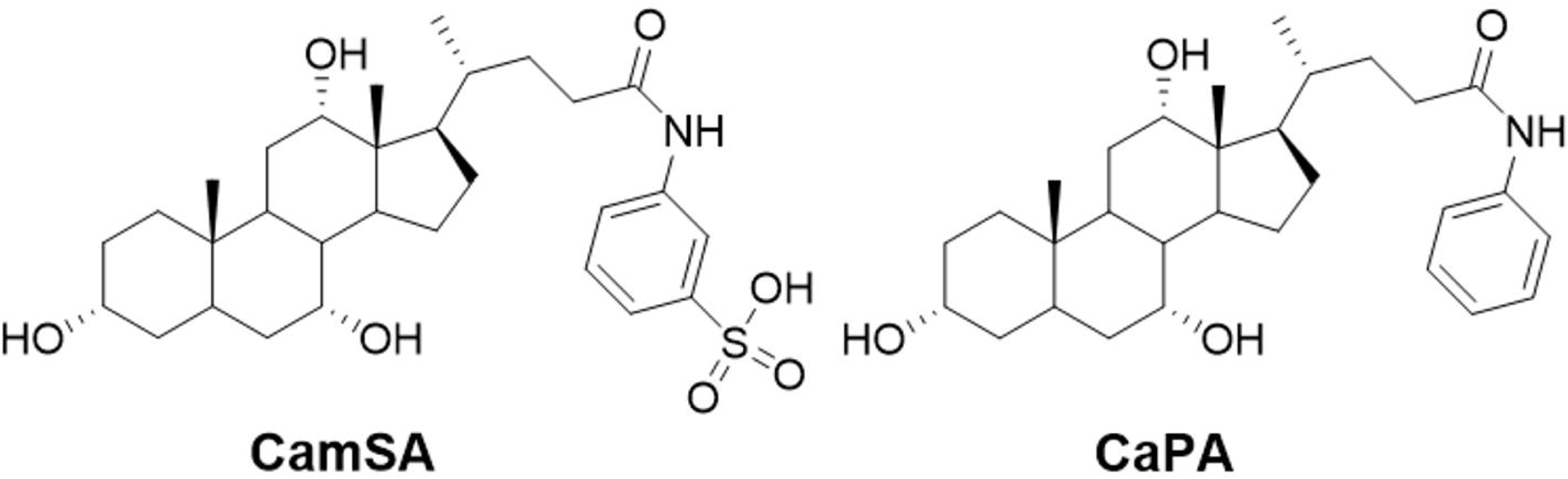
Structure of bile salt anti-germinants used in this study. CamSA is a cholan-24-amides containing a *m*-sulfanilic acid sidechain. CaPA is a cholan-24-amides containing an aniline sidechain.

In this study, we found that CDI prophylaxis showed clear sexual dimorphism. Even though male mice developed less severe CDI, they were also more refractory to treatment. On the other hand, anti-germinants protected female mice from developing CDI during most stages of their estrous cycle. Interestingly, infection sexual dimorphism was reversed in hamsters, with male hamsters developing more severe CDI signs than females. Furthermore, anti-germinant compounds protected female hamsters more strongly than male mice.

## Materials and Methods

### Materials

*C. difficile* strains R20291 and 9001966 were donated by Professor Nigel Minton at the University of Nottingham, United Kingdom. Synthetic bile salt analogs were prepared by Professor Steven M. Firestine at Wayne State University, Michigan, All growth of *C. difficile* strains was done in a Coy Laboratories vinyl anaerobic chamber (5% H_2_, 5% CO_2_, 95% N_2_).

### Animals

All procedures involving animals in this study were performed in accordance with the Guide for Care and Use of Laboratory Animals outlined by the National Institutes of Health. The protocol was reviewed and approved by the Institutional Animal Care and Use Committee (IACUC) at the University of Nevada, Las Vegas (Permit Number: R0914-297). Weaned C57BL/6 mice and weaned Golden Syrian hamsters were purchased from Charles River Laboratories (Wilmington, MA, USA). Mice were housed in groups of five per cage and hamsters were housed individually at the University of Nevada, Las Vegas animal care facility. Upon arrival at the facility, animals were allowed to acclimate for at least one week prior to the start of experimentation. All bedding, cages, food, and water were autoclaved prior to contact with animals. All post-challenge manipulations were performed within a biosafety level 2 laminar flow hood.

#### C. difficile spore harvest and purification

*C. difficile* cells from frozen stocks were streak plated in an anaerobic chamber onto BHI agar supplemented with 2% yeast extract, 0.1% L-cysteine-HCl, and 0.05% sodium taurocholate (BHIS) to yield colonies (15). After 48 hours, a single colony was inoculated into BHI broth supplemented with 0.5% yeast extract and incubated for 48 hours. The inoculated broth was then spread plated onto multiple BHI agar plates prepared as described above. Inoculated plates were incubated for 7 days at 37 °C.

Plates were then flooded with ice-cold deionized (DI) water. Cells and spores were harvested by scraping bacterial colonies from the plates. The cell and spore mixtures were pelleted by centrifugation at 8,000 ×g for 5 minutes, resuspended in DI water, and pelleted again. This washing step was repeated twice more. The cell/spore mixture was then centrifuged through a 20% to 50% HistoDenz gradient at 18,200 ×g for 30 minutes with no brake (22). Under these conditions, spores pelleted at the bottom of the centrifuge tube, while cell debris remained above the 20% HistoDenz layer. Pelleted spores were transferred to a clean centrifuge tube and washed five times before storing in DI water at 4 °C. To determine spore purity, selected samples were stained using the Schaeffer-Fulton endospore staining method (23) or were visualized by phase-contrast microscopy. Spore preparations used were greater than 95% pure.

### Estrous cycle tracking in mice

The estrous cycle tracking method performed in this study was adapted from *McLean et al., 2012* [34]. Mice estrous stages were tracked for at least one week prior to any manipulation to make sure animals were cycling normally. Animals were tracked daily for their estrous cycle stage by vaginal lavage at the same time each morning throughout the duration of the study. Briefly, excess urine is dabbed off using Kimwipes low-lint wipes. A micropipette was used to lavage 20 μL of DI water into the vaginal orifice of the animal. The vaginal orifice was flushed 3-4 times using the micropipette until the suspension appeared mildly cloudy. The 20 μL sample was then transferred onto a clean glass microscope slide and left to air dry for at least 1 hour. Animals were monitored to ensure there were no signs of vaginal infection.

Air-dried vaginal sample slides were dipped into a 2% crystal violet solution inside a Coplin jar for 1 minute. Slides were then moved to a Coplin jar containing fresh DI water to wash off excess stain. Slides were then dabbed gently with clean wipes, allowed to air dry, and examined using a compound light microscope at 100x and 400x magnification.

Using vaginal cytology and staining, estrous cycle stages can be identified based on the presence of either nucleated epithelial cells, cornified (keratinized) anucleated epithelial cells, leukocytes, or combinations of these cell types (**Fig. S1**). The proestrus stage is dominated by the presence of nucleated epithelial cells (blue arrows). In contrast, when animals are in the estrus stage, cornified anucleated epithelial cells are in abundance (yellow arrows). This stage can also show the presence of small specks of bacteria adhering to the epithelial cells (black arrow in the insert). In the metestrus stage, all three major cell types are present. When animals transition to the diestrus stage, samples contain mostly leukocytes (gray arrows) with scattered appearances of a few nucleated epithelial cells.

Cytologic characteristics can also show intermediate stages of the estrous cycle (Fig. S1) [53]. Small, irregular shaped cells (green arrows) can signify the formation of nucleated cells and are representative of late diestrus/early proestrus. Some lingering leukocytes may also be present at this stage. Late proestrus/early estrus is usually marked by a transition of nucleated cells to enucleation leaving a halo-like appearance (pink arrow) in the middle of the cell.

### Murine CDI Model

The murine CDI model used in this study was adapted from Chen *et al.* [54]. Mice were given three consecutive days of antibiotic cocktail containing kanamycin (0.4 mg/ml), gentamicin (0.035 mg/ml), colistin (850 U/ml), metronidazole (0.215 mg/ml), and vancomycin (0.045 mg/ml) *ad libitum* (14). Mice were then given autoclaved DI water for the remainder of the experiment. On the day prior to *C. difficile* challenge (day -1), mice were given an intraperitoneal (IP) injection of 10 mg/kg clindamycin. On the day of infection (day 0), experimental groups were challenged with 10^8^ *C. difficile* spores by oral gavage. To test for CDI prophylaxis, mice were given 50 mg/kg daily gavage doses of either CamSA or CaPA dissolved in DMSO at 0-, 24-, and 48-hours post-challenge. As control, a group of animals were given neat DMSO excipient.

Mice were observed for signs of CDI twice daily and disease severity was scored according to a CDI sign rubric adapted from previously published work [55]. According to the rubric, pink anogenital area, mild wet tail, and weight loss of 8-15% were each given an individual score of 1. Red anogenital area, lethargy/distress, increased diarrhea/soiled bedding, hunched posture, and weight loss greater than 15% were each given an individual score of 2. After adding all scores, animals scoring less than 3 were indistinguishable from noninfected controls and were considered non-diseased. Animals scoring 3–4 were considered to have mild CDI. Animals scoring 5–6 were considered to have moderate CDI. Animals scoring greater than 6 were considered to have severe CDI and were immediately culled.

### Hamster CDI Model

The hamster CDI model used in this study was adapted from procedures reported previously by *Howerton, et al.* [56]. Hamsters were orogastrically dosed with 30 mg/kg clindamycin 1 day prior to infection (day -1). On day 0, animals were challenged with 50 *C. difficile* strain 630 spores. Hamsters were also given a subclinical dose of 1 mg/kg vancomycin daily from day 0 to day 2 to prevent clindamycin-induced colitis. To test for CDI prophylaxis, hamsters were given a 300 mg/kg daily gavage dose of CaPA dissolved in DMSO for 11 days beginning on the day of infection. Animals were observed for CDI signs. Symptomatic animals were immediately culled.

### Statistical Analyses

For the murine CDI model, severity of signs was analyzed by box-and-whisker plots with a minimum of three independent values (n ≥ 3). Single factor ANOVA or one-tailed *t*-test were performed using *R* ((3.5.0) to assess differences between groups at every time point. Post-hoc comparisons of significant ANOVAs were calculated using the Scheffe test for pairwise comparison among all groups. For the hamster model, statistical differences in Kaplan-Meier survival plots were performed using the log-rank test.

## Results

### Sexual dimorphism of mouse response to anti-germinant prophylactics

We had previously shown that male mice infected with spores from *C. difficile* strains 9001966 or R20291 developed less severe CDI than their female counterparts [57]. To evaluate the effect of sexual dimorphism in CDI prevention, male mice (n=5 per group) were challenged with *C. difficile* strain 9001966 spores and treated with were treated with either DMSO excipient (**Fig. 2A**, solid blue columns), CamSA (dashed blue columns), or CaPA (checkered blue columns). (**Fig. 2A**). Even though untreated male mice developed less severe CDI than untreated female mice (**Fig. 2B**), males did not respond to the CDI prophylaxis activity of either CamSA or CaPA at any time during the infection process (**Fig. 2A**).

**Figure 2.**
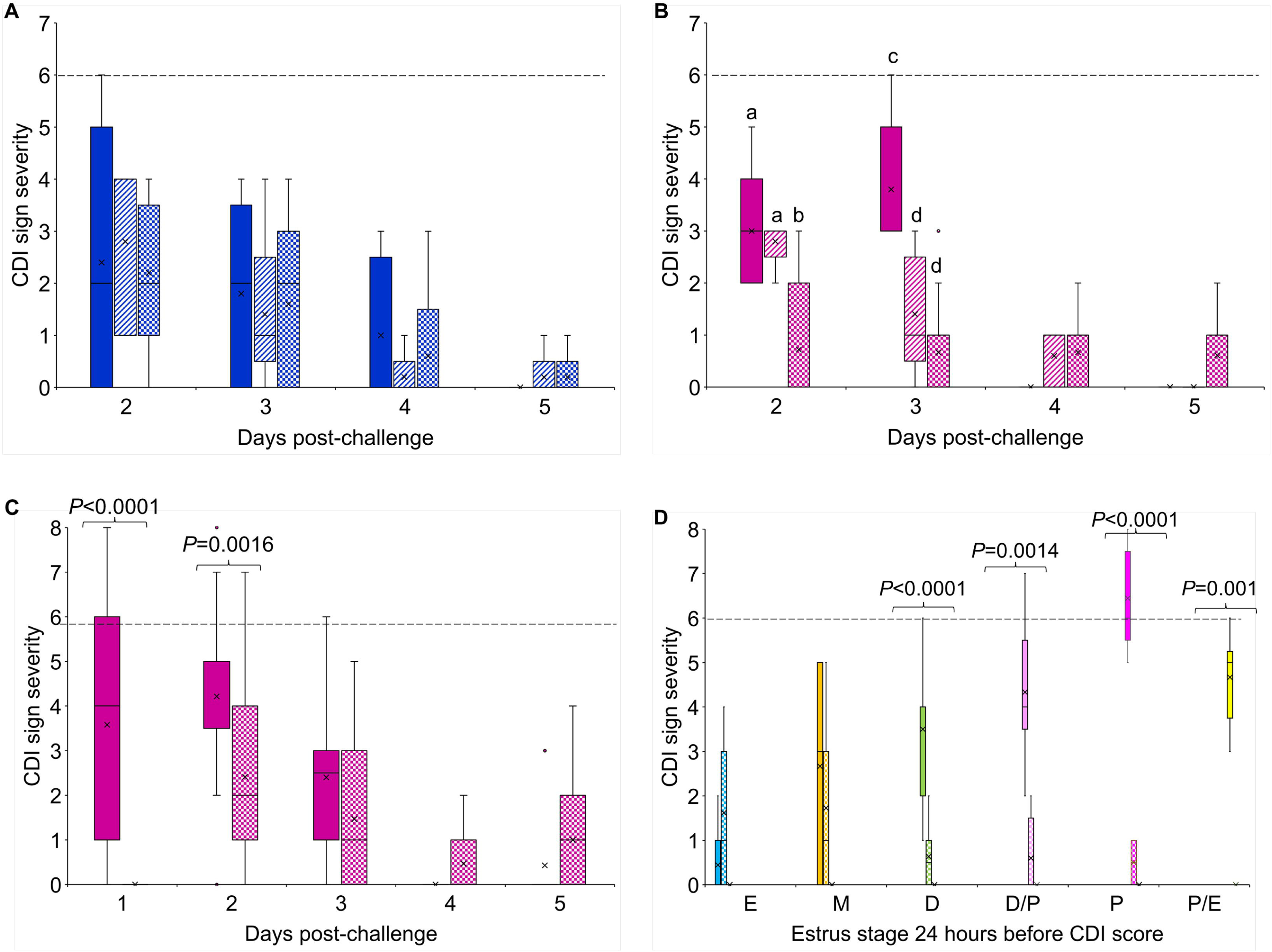
Effect of sex and estrous cycle on murine CDI prophylaxis. (A) Box-and-whisker plots of CDI sign severity in male mice challenged with *C. difficile* spores from strain 9001966 and treated with DMSO (blue solid bars), CamSA (blue striped bars), or CaPA (blue checkered bars). (B) Box-and-whisker plots of CDI sign severity in female mice challenged with *C. difficile* spores from strain 9001966 and treated with DMSO (pink solid bars), CamSA (pink striped bars), or CaPA (pink checkered bars). (C) Box-and-whisker plots of CDI sign severity in female mice challenged with *C. difficile* spores from strain R20291 and treated with DMSO (pink solid bars), or CaPA (pink checkered bars). (D) Cumulative box-and-whisker plots of CDI sign severity caused by *C. difficile* strain R20291 plotted against their prior-day estrous stages and treated with DMSO (solid bars) or with CaPA (checkered bars). Animals that were at estrus (blue bars), metestrus (orange bars), diestrus (green bars), late diestrus/early proestrus (pink bars), proestrus (magenta bars), and late proestrus/early estrus (yellow bars) at any day were respectively pooled together. CDI sign severity and estrous cycle stages were determined daily. Signs of severity were determined based on the CDI scoring rubric reported previously [55]. For panels A and B, single-factor ANOVA was performed at every time point to assess daily differences among the sign severity means of the DMSO-treated, CaPA-treated, and CamSA-treated groups. ANOVA results with *P* values of <0.05 were analyzed *post hoc* using the Scheffe test for pairwise comparison between daily groups. Columns that are labeled with different letters are statistically different. For panels C and D, daily statistical differences between DMSO-treated and CamSA-treated animals were determined by one-tailed *t*-test.

In contrast, CaPA-treated female mice (**Fig. 2B**, checkered pink columns) showed statistically less severe CDI symptoms on both days 2 and 3 post-challenged compared to untreated controls (solid pink columns). CamSA treatment of female mice was less effective and showed protection only on day 3 post-infection (dashed pink columns).

CDI prophylaxis by CaPA in female mice (n=10 per group) was also evaluated against the more virulent *C. difficile* strain R20291 (**Fig. 2C**). Daily pairwise comparison shows that CaPA-treated female mice (checkered pink columns) developed less severe CDI symptoms at days 1 and 2 post-infection compared to control DMSO-treated animals (solid pink columns). Because animals infected with *C. difficile* strain R20291 showed more virulent CDI signs, we used this hypervirulent strain for all subsequent experiments.

### Effect of estrous stage on CDI prophylaxis

We have previously shown that the estrus stage on the day of infection affect CDI severity during infection progression [57]. Indeed, animals challenged with spores while on estrus were quite resistant to CDI. In contrast, animals challenged while on proestrus stage developed severe CDI.

To understand the effect of estrous cycle on CDI prophylaxis, female mice at different stages of the estrous cycle were infected with hypervirulent *C. difficile* strain R20291 (**Fig. 2D**) and treated with either DMSO excipient (n=60, solid-colored columns) or CaPA (n=18, checkered-colored columns). Pairwise comparisons between DMSO-treated animals and CaPA treated animals showed that animals that were at any time in estrus (blue columns) or metestrus (orange columns) when infected failed to respond to CaPA-treatment. In contrast, CaPA-treatment resulted in statistically significant reduction of next-day CDI sign severity for animals that were at any point in diestrus (green columns), diestrus/proestrus (pink columns), proestrus (magenta columns), and proestrus/estrus (yellow columns).

### Sexual dimorphism of hamster response to anti-germinant prophylactics

Sexual dimorphism of CDI severity was also evaluated in the hamster model of CDI (n=5 per group). In contrast to mice, male hamsters showed slightly higher, but not statistically significant, mortality than females (**Fig. 3**, p=0.06). Indeed, all male hamsters were deceased by day 3 (solid blue circles), while 40% of female hamsters survived to the end of the experiment (solid pink circles). Moreover, when hamsters were treated with CaPA, male hamster mortality was delayed, but all animals became moribund by day 6 (checkered blue squares), whereas female hamsters treated with CaPA (checkered pink circles) showed statistically higher survival rate than CaPA-treated males (p=0.03).

**Figure 3.**
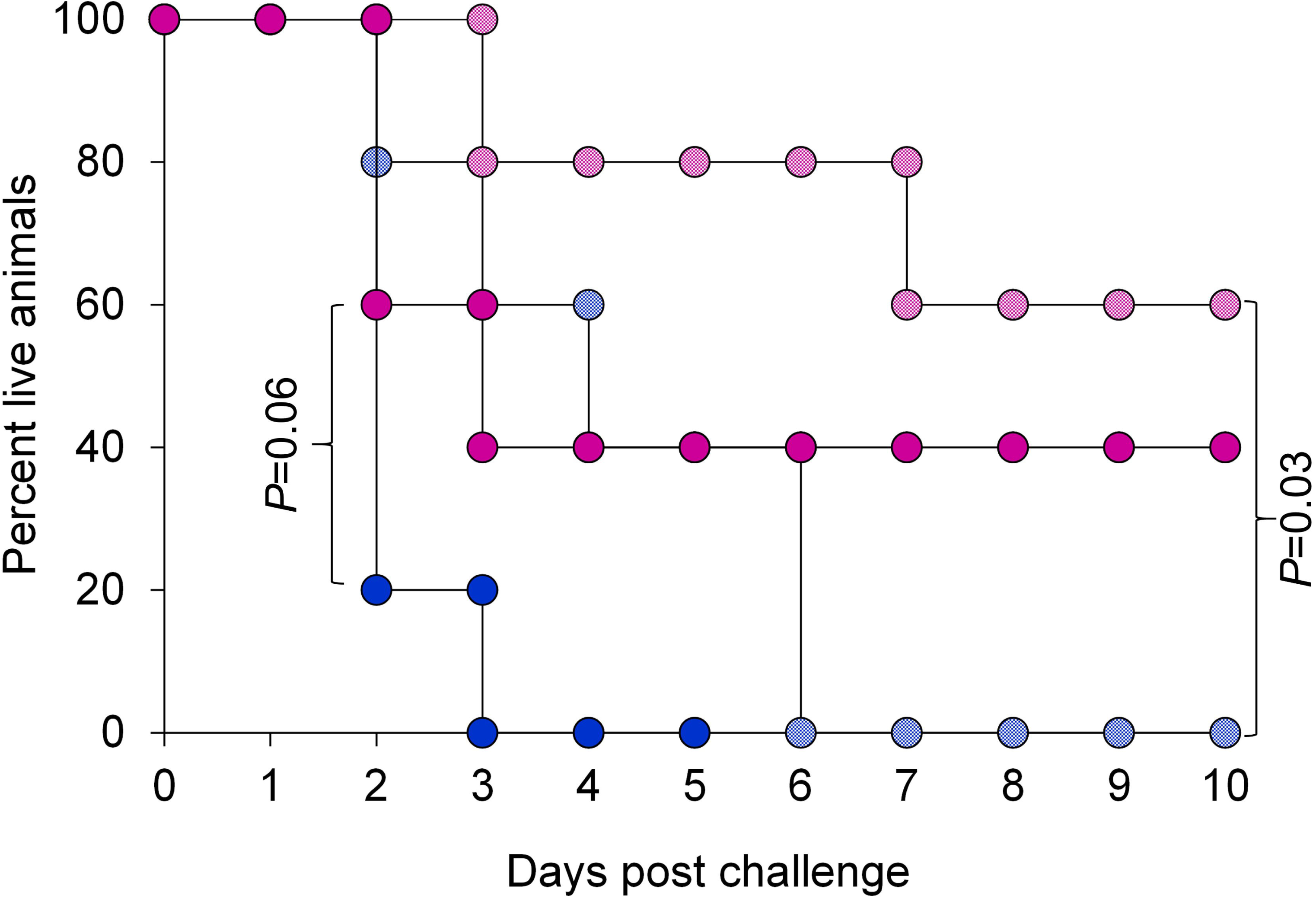
Effect of sex on hamster CDI survival. Kaplan-Meier survival plot for DMSO-treated female hamsters (solid pink circles), CaPA-treated female hamsters (checkered pink circles), DMSO-treated male hamsters (solid blue circles), and CaPA-treated male hamsters (checkered blue circles) challenged with *C. difficile* spores from strain R20291. Statistical survival comparisons between groups were performed by log rank test.

### Discussion and Conclusions

Anti-germination approaches represent a new therapeutic venue to prevent CDI onset. When evaluated against the slow onset *C. difficile* strain 9001966, we found sexual dimorphism in CDI prophylaxis. Indeed, even though males developed less severe CDI than females, anti-germination compounds did not affect the course of male infection. In contrast, female mice were protected from the most severe CDI signs by both CaPA and CamSA, although CaPA seems to be more effective over time.

CaPA was also active in preventing CDI in female mice infected with hypervirulent strain R20291. Intriguingly, the effect of CaPA on CDI sign severity shows the same 24-hour delay following each estrous stage. The only exception to CaPA prophylaxis activity were animals that came out of estrus and metestrus. The estrous stage independence from CaPA action is easily explainable since following estrus, animals naturally do not develop CDI, making CaPA treatment redundant. The refractivity to CaPA of animals coming from metestrus is baffling, since in all other estrous stages, CaPA was able to completely prevent CDI onset. The metestrus abnormality in the CDI prevention/estrous cycle axis needs to be investigated further.

Rodent models of CDI have been invaluable to study infection progression as well as to test for new treatments [58, 59]. Mice are quite resistant to CDI and require pre-conditioning with aggressive antibiotic treatment [54]. Nevertheless, once CDI is established, the murine model shows symptom progression that mimics human CDI [59].

Even though the hamster is the most used CDI model, disease onset is fulminant [60]. These conditions are rarely seen in human primary CDI. On the other hand, vancomycin treatment of infected hamsters results in delayed sign onset similar to human CDI relapse [60, 61]. Hence, the two CDI rodent models complement each other and cover the spectrum of human CDI symptomatology [56, 59].

Because CDI onset and symptomatology diverge in mouse and hamsters, it is not uncommon to find distinct responses to infection and treatments. Indeed, we recently found that carbohydrate-rich diets are protective in murine CDI [55], but result in delayed CDI onset in hamsters [62]. Here we found a similar mirror effect between the two models: while female mice developed more severe CDI signs, female hamsters were slightly more resistant to CDI-mediated mortality and responded better to CDI prophylaxis. The causes for these interspecies differences are currently under investigation.

Together, these data suggest that CDI prophylaxis is modulated by sexual dimorphism. In both rodent models, female mice were more responsive to anti-germinant compounds and were better protected from CDI. Furthermore, CDI protection was dependent on the estrous phase at which female mice were infected.

## Acknowledgments

This work was supported by the National Institute of Health [grant numbers R01-AI109139 and R16AI175022]. The authors thank Prof. Steve Firestine from Wayne State University for the synthesis of CaPA, and Prof. Nigel Minton from University of Nottingham for providing clinical *C. difficile* strains.

## References

1. Dicks LMT, Mikkelsen LS, Brandsborg E, Marcotte H. *Clostridium difficile*, the Difficult “Kloster” Fuelled by Antibiotics. Current Microbiology. 2019;76(6):774–82; doi: 10.1007/s00284-018-1543-8.

2. CDC: Antibiotic Resistance Threats in the United States 2019. In. Edited by Services USDoHaH. Atlanta, GA; 2019.

3. Zhang S, Palazuelos-Munoz S, Balsells EM, Nair H, Chit A, Kyaw MH. Cost of hospital management of Clostridium difficile infection in United States—a meta-analysis and modelling study. J BMC Infectious Diseases. 2016;16(1):447; doi: 10.1186/s12879-016-1786-6.

4. Ghose C. *Clostridium difficile* Infection in the Twenty-First Century. Emerging microbes & Infections. 2013;2(9):e62–e; doi: 10.1038/emi.2013.62.

5. Olsen MA, Young-Xu Y, Stwalley D, Kelly CP, Gerding DN, Saeed MJ, et al. The Burden of *Clostridium difficile* Infection: Estimates of the Incidence of CDI from U.S. Administrative Databases. BMC Infect Dis. 2016;16:177–; doi: 10.1186/s12879-016-1501-7.

6. McDonald LC, Gerding DN, Johnson S, Bakken JS, Carroll KC, Coffin SE, et al. Clinical Practice Guidelines for Clostridium difficile Infection in Adults and Children: 2017 Update by the Infectious Diseases Society of America (IDSA) and Society for Healthcare Epidemiology of America (SHEA). Clinical Infectious Diseases. 2018;66(7):e1–e48; doi: 10.1093/cid/cix1085.

7. Mylonakis E, Ryan ET, Calderwood SB. *Clostridium difficile*-Associated Diarrhea: A Review. Arch Intern Med. 2001;161(4):525–33.

8. Greig J. Fidaxomicin in the Treatment of Clostridium difficile-Associated Diarrhoea. Clinical Drug Investigation. 2013;33(1):93–4; doi: 10.1007/s40261-012-0044-y.

9. Bassetti M, Villa G, Pecori D, Arzese A, Wilcox M. Epidemiology, diagnosis and treatment of Clostridium difficile infection. Expert Review of Anti-infective Therapy. 2012;10(12):1405–23; doi: 10.1586/eri.12.135.

10. Leuzzi R, Adamo R, Scarselli M. Vaccines against Clostridium difficile. Hum Vaccin Immunother. 2014;10(6):1466–77; doi: 10.4161/hv.28428.

11. Yip C, Phan J, Abel-Santos E. Treatment of *Clostridium difficile* Infections. In: Firestine SM, Lister T, editors. Antibiotic Drug Discovery: New Targets and Molecular Entities. London, UK: The Royal Society of Chemistry; 2017. p. 1–19

12. Bakken JS, Borody T, Brandt LJ, Brill JV, Demarco DC, Franzos MA, et al. Treating Clostridium difficile Infection With Fecal Microbiota Transplantation. Clinical Gastroenterology and Hepatology. 2011;9(12):1044–9; doi: 10.1016/j.cgh.2011.08.014.

13. Kuijper EJ, Allegretii J, Hawkey P, Sokol H, Goldenberg S, Ianiro G, et al. A necessary discussion after transmission of multidrug-resistant organisms through faecal microbiota transplantations. The Lancet Infectious Diseases. 2019;19(11):1161–2; doi: 10.1016/S1473-3099(19)30545-6.

14. Eze P, Balsells E, Kyaw MH, Nair H. Risk factors for Clostridium difficile infections - an overview of the evidence base and challenges in data synthesis. Journal of global health. 2017;7(1):010417–; doi: 10.7189/jogh.07.010417.

15. Freeman J, Wilcox MH. Antibiotics and Clostridium difficile. Microbes and Infection. 1999;1(5):377–84; doi: 10.1016/S1286-4579(99)80054-9.

16. Bobak D, Arfons LM, Creger RJ, Lazarus HM. Clostridium difficile-associated disease in human stem cell transplant recipients: coming epidemic or false alarm[quest]. Bone Marrow Transplant. 2008;42(11):705–13.

17. Arslan H, Inci EK, Azap OK, Karakayali H, Torgay A, Haberal M. Etiologic agents of diarrhea in solid organ recipients. Transplant Infectious Disease. 2007;9(4):270–5.

18. Sanchez Travis H, Brooks John T, Sullivan PS, Juhasz M, Mintz E, Dworkin Mark S, et al. Bacterial Diarrhea in Persons with HIV Infection, United States, 1992-2002. Clinical Infectious Diseases. 2005;41(11):1621–7; doi: doi:10.1086/498027.

19. Vázquez-Martínez ER, García-Gómez E, Camacho-Arroyo I, González-Pedrajo B. Sexual dimorphism in bacterial infections. Biology of Sex Differences. 2018;9(1):27; doi: 10.1186/s13293-018-0187-5.

20. Lucado J GCEA: Clostridium difficile Infections (CDI) in Hospital Stays, 2009. In. Edited by (US) AfHRaQ. Rockville (MD); 2012.

21. Elixhauser A SC, Gould C. : Readmissions following Hospitalizations with Clostridium difficile Infections, 2009. In. Edited by (US) AfHRaQ. Rockville (MD); 2012.

22. Saffouri G, Gupta A, Loftus EV, Baddour LM, Pardi DS, Khanna S. The incidence and outcomes from Clostridium difficile infection in hospitalized adults with inflammatory bowel disease. Scandinavian Journal of Gastroenterology. 2017;52(11):1240–7; doi: 10.1080/00365521.2017.1362466.

23. Choi H-Y, Park S-Y, Kim Y-A, Yoon T-Y, Choi J-M, Choe B-K, et al. The epidemiology and economic burden of Clostridium difficile infection in Korea. BioMed research international. 2015;2015:510386–; doi: 10.1155/2015/510386.

24. King A, Mullish BH, Williams HRT, Aylin P. Comparative epidemiology of Clostridium difficile infection: England and the USA. International Journal for Quality in Health Care. 2017;29(6):785–91; doi: 10.1093/intqhc/mzx120.

25. Garey KW, Dao-Tran TK, Jiang ZD, Price MP, Gentry LO, DuPont HL. A clinical risk index for Clostridium difficile infection in hospitalised patients receiving broad-spectrum antibiotics. Journal of Hospital Infection. 2008;70(2):142–7; doi: 10.1016/j.jhin.2008.06.026.

26. Johnson S. Recurrent Clostridium difficile infection: A review of risk factors, treatments, and outcomes. Journal of Infection. 2009;58(6):403–10; doi: 10.1016/j.jinf.2009.03.010.

27. Garey KW, Sethi S, Yadav Y, DuPont HL. Meta-analysis to assess risk factors for recurrent Clostridium difficile infection. Journal of Hospital Infection. 2008;70(4):298– 304; doi: 10.1016/j.jhin.2008.08.012.

28. Rouphael NG, O’Donnell JA, Bhatnagar J, Lewis F, Polgreen PM, Beekmann S, et al. Clostridium difficile–associated diarrhea: an emerging threat to pregnant women. American Journal of Obstetrics and Gynecology. 2008;198(6):635.e1–.e6; doi: 10.1016/j.ajog.2008.01.062.

29. Cózar-Llistó A, Ramos-Martinez A, Cobo J. Clostridium difficile Infection in Special High-Risk Populations. Infectious diseases and therapy. 2016;5(3):253–69; doi: 10.1007/s40121-016-0124-z.

30. Meda M, Virgincar N, Gentry V, Walker A, Macdonald N, Hooper M, et al. Clostridium difficile infection in pregnant and postpartum women in 2 hospitals and a review of literature. American Journal of Infection Control. 2019;47(1):e7–e14; doi: 10.1016/j.ajic.2018.06.001.

31. Garey KW, Jiang Z-D, Yadav Y, Mullins B, Wong K, Dupont HL. Peripartum Clostridium difficile infection: case series and review of the literature. American Journal of Obstetrics and Gynecology. 2008;199(4):332–7; doi: 10.1016/j.ajog.2008.05.001.

32. Draper CF, Duisters K, Weger B, Chakrabarti A, Harms AC, Brennan L, et al. Menstrual cycle rhythmicity: metabolic patterns in healthy women. Sci Rep. 2018;8(1):14568; doi: 10.1038/s41598-018-32647-0.

33. Byers SL, Wiles MV, Dunn SL, Taft RA. Mouse Estrous Cycle Identification Tool and Images. PLOS ONE. 2012;7(4):e35538; doi: 10.1371/journal.pone.0035538.

34. McLean AC, Valenzuela N, Fai S, Bennett SAL. Performing Vaginal Lavage, Crystal Violet Staining, and Vaginal Cytological Evaluation for Mouse Estrous Cycle Staging Identification. JoVE. 2012;(67):e4389; doi: doi:10.3791/4389.

35. Burns DA, Heap JT, Minton NP. *Clostridium difficile* Spore Germination: an Update. Research in Microbiology. 2010;161(9):730–4; doi: 10.1016/j.resmic.2010.09.007.

36. Giel JL, Sorg JA, Sonenshein AL, Zhu J. Metabolism of Bile Salts in Mice Influences Spore Germination in *Clostridium difficile*. PLOS ONE. 2010;5(1):e8740; doi: 10.1371/journal.pone.0008740.

37. Francis MB, Allen CA, Shrestha R, Sorg JA. Bile acid recognition by the *Clostridium difficile* germinant receptor, Cspc, is important for establishing infection. PLOS Pathogens. 2013;9(5):e1003356; doi: 10.1371/journal.ppat.1003356.

38. Sorg JA, Sonenshein AL. Chenodeoxycholate is an inhibitor of *Clostridium difficile* spore germination. J Bacteriol. 2009;191(3):1115–7; doi: 10.1128/jb.01260-08.

39. Sorg JA, Sonenshein AL. Bile salts and glycine as cogerminants for *Clostridium difficile* spores. J Bacteriol. 2008;190(7):2505–12; doi: 10.1128/jb.01765-07.

40. Howerton A, Ramirez N, Abel-Santos E. Mapping interactions between germinants and *Clostridium difficile* spores. J Bacteriol. 2011;193(1):274–82; doi: 10.1128/jb.00980-10.

41. Ramirez N, Liggins M, Abel-Santos E. Kinetic evidence for the presence of putative germination receptors in *C. difficile* spores. J Bacteriol. 2010;192(16):4215–22.

42. Alvarez Z, Abel-Santos E. Potential use of inhibitors of bacteria spore germination in the prophylactic treatment of anthrax and Clostridium difficile-associated disease. Expert Review of Anti-infective Therapy. 2007;5(5):783–92; doi: doi:10.1586/14787210.5.5.783.

43. Howerton A, Ramirez N, Abel-Santos E. Mapping Interactions between Germinants and Clostridium difficile Spores. Journal of Bacteriology. 2011;193(1):274–82; doi: 10.1128/jb.00980-10.

44. Liggins M, Ramirez N, Magnuson N, Abel-Santos E. Progesterone Analogs Influence Germination of Clostridium sordellii and Clostridium difficile Spores In Vitro. Journal of Bacteriology. 2011;193(11):2776–83; doi: 10.1128/jb.00058-11.

45. Sorg JA, Sonenshein AL. Chenodeoxycholate is an inhibitor of *Clostridium difficile* spore germination. Journal of Bacteriology. 2009;191(3):1115–7; doi: 10.1128/jb.01260-08.

46. Yip C, Okada NC, Howerton A, Amei A, Abel-Santos E. Pharmacokinetics of CamSA, a potential prophylactic compound against Clostridioides difficile infections. Biochemical Pharmacology. 2021;183:114314; doi: 10.1016/j.bcp.2020.114314.

47. Phan JR, Do DM, Truong MC, Ngo C, Phan JH, Sharma SK, et al. An Aniline-Substituted Bile Salt Analog Protects both Mice and Hamsters from Multiple Clostridioides difficile Strains. Antimicrobial Agents and Chemotherapy. 2022;66(1):e01435–21; doi: doi:10.1128/AAC.01435-21.

48. Sharma SK, Yip C, Esposito EX, Sharma PV, Simon MP, Abel-Santos E, et al. The Design, Synthesis, and Characterizations of Spore Germination Inhibitors Effective against an Epidemic Strain of Clostridium difficile. Journal of medicinal chemistry. 2018;61(15):6759–78; doi: 10.1021/acs.jmedchem.8b00632.

49. Howerton A, Seymour CO, Murugapiran SK, Liao Z, Phan JR, Estrada A, et al. Effect of the Synthetic Bile Salt Analog CamSA on the Hamster Model of Clostridium difficile Infection. Antimicrobial Agents and Chemotherapy. 2018;62(10).

50. Howerton A, Patra M, Abel-Santos E. Fate of ingested Clostridium difficile spores in mice. PloS one. 2013;8(8):e72620–e; doi: 10.1371/journal.pone.0072620.

51. Howerton A, Patra M, Abel-Santos E. A New Strategy for the Prevention of Clostridium difficile Infection. Journal of Infectious Diseases. 2013;207(10):1498–504; doi: 10.1093/infdis/jit068.

52. Sattar A, Thommes P, Payne L, Warn P, Vickers RJ. SMT19969 for *Clostridium difficile* infection (CDI): In Vivo Efficacy Compared with Fidaxomicin and Vancomycin in the Hamster Model of CDI. The Journal of antimicrobial chemotherapy. 2015;70(6):1757– 62; doi: 10.1093/jac/dkv005.

53. Cora MC, Kooistra L, Travlos G. Vaginal Cytology of the Laboratory Rat and Mouse: Review and Criteria for the Staging of the Estrous Cycle Using Stained Vaginal Smears. Toxicol Pathol. 2015;43(6):776–93; doi: 10.1177/0192623315570339.

54. Chen X, Katchar K, Goldsmith JD, Nanthakumar N, Cheknis A, Gerding DN, et al. A Mouse Model of Clostridium difficile-Associated Disease. Gastroenterology. 2008;135(6):1984–92.

55. Mefferd CC, Bhute SS, Phan JR, Villarama JV, Do DM, Alarcia S, et al. A High-Fat/High-Protein, Atkins-Type Diet Exacerbates *Clostridioides (Clostridium) difficile* Infection in Mice, whereas a High-Carbohydrate Diet Protects. mSystems. 2020;5(1):e00765–19; doi: 10.1128/mSystems.00765-19.

56. Howerton A, Seymour CO, Murugapiran SK, Liao Z, Phan JR, Estrada A, et al. Effect of the Synthetic Bile Salt Analog CamSA on the Hamster Model of *Clostridium difficile* Infection. Antimicrobial Agents and Chemotherapy. 2018;62(10): e02251–17.

57. Phan JR, Washington M, Do DM, Mata TV, Niamba M, Heredia E, et al. Effects of sexual dimorphism and estrous cycle on C. difficile infections in rodent models. bioRxiv. 2023:2023.07.05.547871; doi: 10.1101/2023.07.05.547871.

58. Warn P, Thommes P, Sattar A, Corbett D, Flattery A, Zhang Z, et al. Disease Progression and Resolution in Rodent Models of Clostridium difficile Infection and Impact of Antitoxin Antibodies and Vancomycin. 2016;60(11):6471–82; doi: 10.1128/AAC.00974-16 %J Antimicrobial Agents and Chemotherapy.

59. Best EL, Freeman J, Wilcox MH. Models for the study of *Clostridium difficile* infection. Gut Microbes. 2012 3(2):145–67.

60. Mullany P, Roberts A, Douce G, Goulding D. Refinement of the Hamster Model of Clostridium difficile Disease. In: Clostridium difficile. Humana Press; 2010. p. 215–27.

61. Anton PM, O’Brien M, Kokkotou E, Eisenstein B, Michaelis A, Rothstein D, et al. Rifalazil Treats and Prevents Relapse of Clostridium difficile-Associated Diarrhea in Hamsters. Antimicrobial Agents and Chemotherapy. 2004;48(10):3975–9; doi: 10.1128/aac.48.10.3975-3979.2004.

62. Bhute SS, Mefferd CC, Phan JR, Ahmed M, Fox-King AE, Alarcia S, et al. A High-Carbohydrate Diet Prolongs Dysbiosis and Clostridioides difficile Carriage and Increases Delayed Mortality in a Hamster Model of Infection. Microbiol Spectr. 2022:e0180421; doi: 10.1128/spectrum.01804-21.

